# Antibiotic action revealed by real-time imaging of the mycobacterial membrane

**DOI:** 10.1101/2022.01.07.475452

**Authors:** Michael G. Wuo, Charles L. Dulberger, Robert A. Brown, Alexander Sturm, Eveline Ultee, Zohar Bloom-Ackermann, Catherine Choi, Ethan C. Garner, Ariane Briegel, Deborah T. Hung, Eric J. Rubin, Laura L. Kiessling

**Author notes:** Corresponding author: Laura L. Kiessling, **Email:**.

## Abstract

The current understanding of mycobacterial cell envelope remodeling in response to antibiotics is limited. Chemical tools that report on phenotypic changes with minimal cell wall perturbation are critical to understanding such time-dependent processes. We employed a fluorogenic chemical probe to image how antibiotics perturb mycobacterial cell envelope assembly in real-time. Time-lapse microscopy revealed that differential antibiotic treatment elicited unique cellular phenotypes, providing a platform for simultaneously monitoring cell envelope construction and remodeling responses. Our data show that rifampicin, which does not directly inhibit cell wall biosynthesis, affords a readily detected mycomembrane phenotype. The fluorogenic probe revealed the production of extracellular vesicles in response to antibiotics, and analyses of these vesicles indicate that antibiotic treatment elicits the release of agents that attenuate macrophage activation.

## Introduction

Mycobacterial cell envelope biogenesis and remodeling are dynamic processes critical to infectivity and recalcitrance to antibiotics (1). Mycolyl arabinogalactan (mAG) is the principal constituent of the cellular envelope recognized by host macrophages (2–4). Changes in envelope architecture play critical roles in colony morphology, pellicle formation, and drug resistance (5–7). Substantial restructuring of the cell envelope (e.g., increased production of mycolic acids, changes in arabinogalactan synthesis, and modifications of peptidoglycan crosslinks) can contribute to bacterial persistence and cellular survival (8– 11). Still, the molecular mechanisms that allow intracellular mycobacteria to survive under antibiotic treatment are enigmatic. Previous studies have evaluated morphological changes in physical dimensions and DNA compaction after antibiotic treatment (Fig 1A) (12). However, these changes alone cannot account for the persistent infections that reestablish following antibiotic treatment. New approaches for evaluating the effects of antibiotics on mycobacterial membrane remodeling can uncover unexpected mechanisms of mycobacterial persistence and new therapeutic strategies that address the threat of antibiotic resistance.

**Figure 1.**
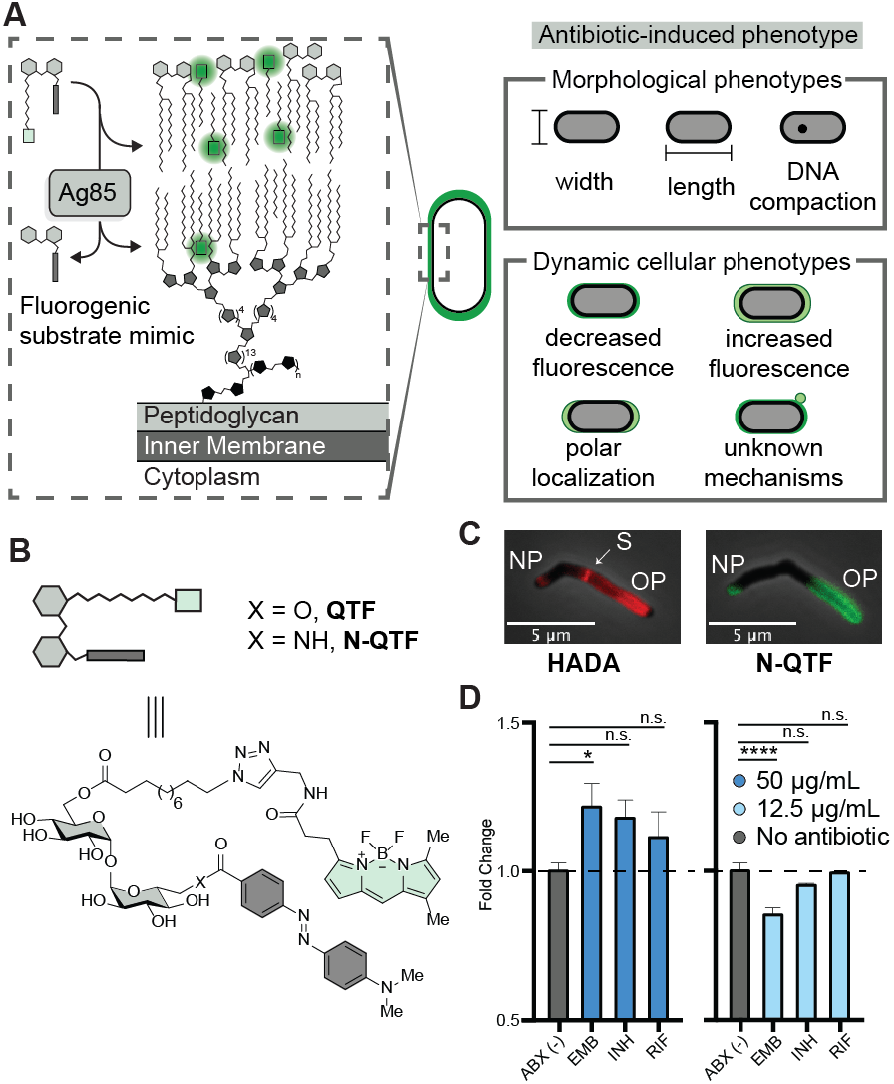
Analysis of N-QTF, a fluorogenic, real-time reporter of mycolyltransferase activity. (A) The fluorogenic substrate (QTF or N-QTF, B) is activated by the Ag85 enzyme complex and serves as a reporter of dynamic antibiotic-induced cellular phenotypes. (B) The fluorogenic probes, QTF and N-QTF, mimic critical features of trehalose monomycolate, the natural substrate of Ag85. (C) Fluorescence labeling of *M. smegmatis* with HADA (left) or N-QTF (right). NP: new pole OP: old pole S: septum (D) Bulk fluorescence measurements show concentration-dependent N-QTF activation differences upon treatment with frontline antibiotics at 50 μg/mL (left, blue) and 12.5 μg/mL (right, light blue). (Ordinary oneway-ANOVA ****p ≤ 0.0001, ***p ≤ 0.001, **p ≤ 0.01, *p ≤0.05 and non-significant >0.05). Data shown are the average of triplicate samples ± SEM in triplicate and representative of two independent experiments.

Mycobacteria are master manipulators of their environment (13). They communicate to host cells through direct mAG contact and secreted molecules. For example, mAG trehalose dimycolate is recognized by pattern recognition receptors on macrophage surfaces (14, 15). Moreover, *Mtb* secrete outer membrane vesicles to influence TLR2-mediated activation of macrophages (16). Many frontline antibiotics act on biosynthetic enzymes involved in mycobacterial cell envelope construction; therefore, strategies that report in real-time on mycomembrane dynamics and secretion mechanisms can illuminate how mycobacterial growth, division, and remodeling influence survival. We identify distinct antibiotic-induced membrane remodeling phenotypes using a fluorogenic reporter of cell envelope construction. Guided by these specific phenotypes, we isolated and characterized antibiotic-induced outer membrane vesicles (OMVs). Unexpectedly, they reduce macrophage activation, suggesting a role in developing rifampicin resistance in *Mtb* infection.

## Results

### N-QTF synthesis and fluorogenic properties

Many fluorescent reporters can report on mycobacterial physiology (17–19); however, many require washing and fixation steps that limit their use for real-time monitoring of mycobacterial growth and division. To address this deficiency, we developed a fluorogenic probe, <u>q</u>uencher trehalose fluorophore (QTF) (20), that reports on the growth of the lipid-rich mycobacterial cell envelope (Fig. 1A). To improve the thermodynamic and hydrolytic stability of the probe, we made a single substitution at the trehalose C6-position to generate an amide-containing derivative, N-quencher trehalose fluorophore or N-QTF (Fig. 1B, Supp. Fig. S1, S2).

We examined N-QTF activation in different mycobacterial species by treating fast-growing *M. smegmatis* (*Msmeg*), or slow-growing *M. marinum* (*Mmar*) *or M. tuberculosis* (*Mtb*) with the fluorogenic probe (5 μM). The emergence of fluorescence over time was commensurate with mycobacterial species doubling time (Supp. Fig. S3). To test the molecular mechanism of N-QTF activation, we subjected N-QTF to mycolyltransferases Ag85A-C and monitored enzymatic activity (Supp. Fig. S3). All mycolyltransferases activated N-QTF; however, Ag85C catalyzed turn-on with the fastest kinetics. Single functional group substitutions can dramatically affect probe processing and fluorescence localization (11). We monitored localization using time-lapse microscopy to determine whether the N-QTF probe exhibits fluorescence at the polar and septal regions of growing mycobacteria as described for QTF (Fig. 1C. Supp Fig. S3). Peptidoglycan labeling with 7-hydroxycoumarin-3-carboxylic acid-3-amino-3-D-alanine at polar regions (Fig. 1C) supports N-QTF enables selective, real-time visualization of envelope biogenesis.

### N-QTF temporal response to antibiotic-treated mycobacteria

Before leveraging N-QTF to examine spatial and temporal fluorogenic phenotypes in response to antibiotic stress, we first determined how antibiotic treatment influences the fluorescence resulting from N-QTF processing. We used a time-course fluorescence assay to monitor *Msmeg* responses to frontline *Mtb* antibiotics ethambutol (EMB), rifampicin (RIF), and isoniazid (INH) at 12.5 μg/mL (low, sub-MIC) and 50 μg/mL (high, super-MIC) concentrations. The bulk fluorescence measured over six hours (two doubling times, Supp Fig. S4) tracked with antibiotic concentration with a higher fluorescent signal at a higher antibiotic concentration (Fig. 1D). A linear correlation of fluorescence and OD600 was observed for all treatment conditions, suggesting that the bacteria could grow and multiply under the assay conditions during this short, early period of observation (Supp. Fig. S4). Of the frontline antibiotics, ethambutol (EMB)-treated cells exhibited the greatest fluorescence increase at high concentrations and most significant fluorescence decrease at low concentrations relative to untreated control. Rifampicin (RIF)-treated cells had only a minor relative fluorescence change, while isoniazid (INH) treatment had an intermediate response. These data suggest that *Msmeg* increases lipid remodeling activity under antibiotic stress and that each anti-mycobacterial agent elicits a distinct cell membrane response.

EMB inhibits the arabinofuranosyltransferases EmbA-C (21). These enzymes generate the cell envelope arabinan layer, to which the lipid-rich mycolic acid layer is appended. Prior studies indicate ethambutol treatment results in an intracellular accumulation and secretion of trehalose mycolates and mycolic acids in the culture media (22, 23). In one study, EMB treatment induced increased expression of the polar scaffolding protein DivIVA (24), a fulcrum for the organization of the polar elongation complex, elongasome. This investigation recorded active growth at the cellular poles; paradoxically, the level of mycolic acid at the cellular poles appeared to be reduced (24). We reasoned that these counterintuitive results likely stem from the non-specific fluorescent dye used to monitor mycolic acid changes. The fluorescence of such dyes depends on the hydrophobicity of the region; therefore, they cannot report on dynamic remodeling.

### Time-lapse microscopy of antibiotic-treated mycobacteria yields specific cellular phenotypes

We postulated that N-QTF could report on lipid restructuring in real-time. Using time-lapse imaging under microfluidic control, we continuously supplied N-QTF to *Msmeg*. We flowed either vehicle or EMB antibiotic (10 μg/mL) into the culture chamber at defined time points (Fig. 2A-C, Movies S1-S4). Changes in fluorescence localization were monitored over time. After one doubling time, EMB-treated *Msmeg* showed bright fluorescence localization to the bacillus’ polar regions. The observed phenotype increased over time, and after 16 hours, we detected ejection of the mycomembrane lipids at these sites. These data suggest that the cell pole is more fluid than previously described, a property that explains why non-specific fluorescent dyes could not report on the lipid changes revealed by N-QTF. These new findings underscore the importance of capturing the dynamics of lipid remodeling under antibiotic treatment, as spatiotemporal resolution provides mechanistic insight into changes in cell wall biosynthetic pathways.

**Figure 2.**
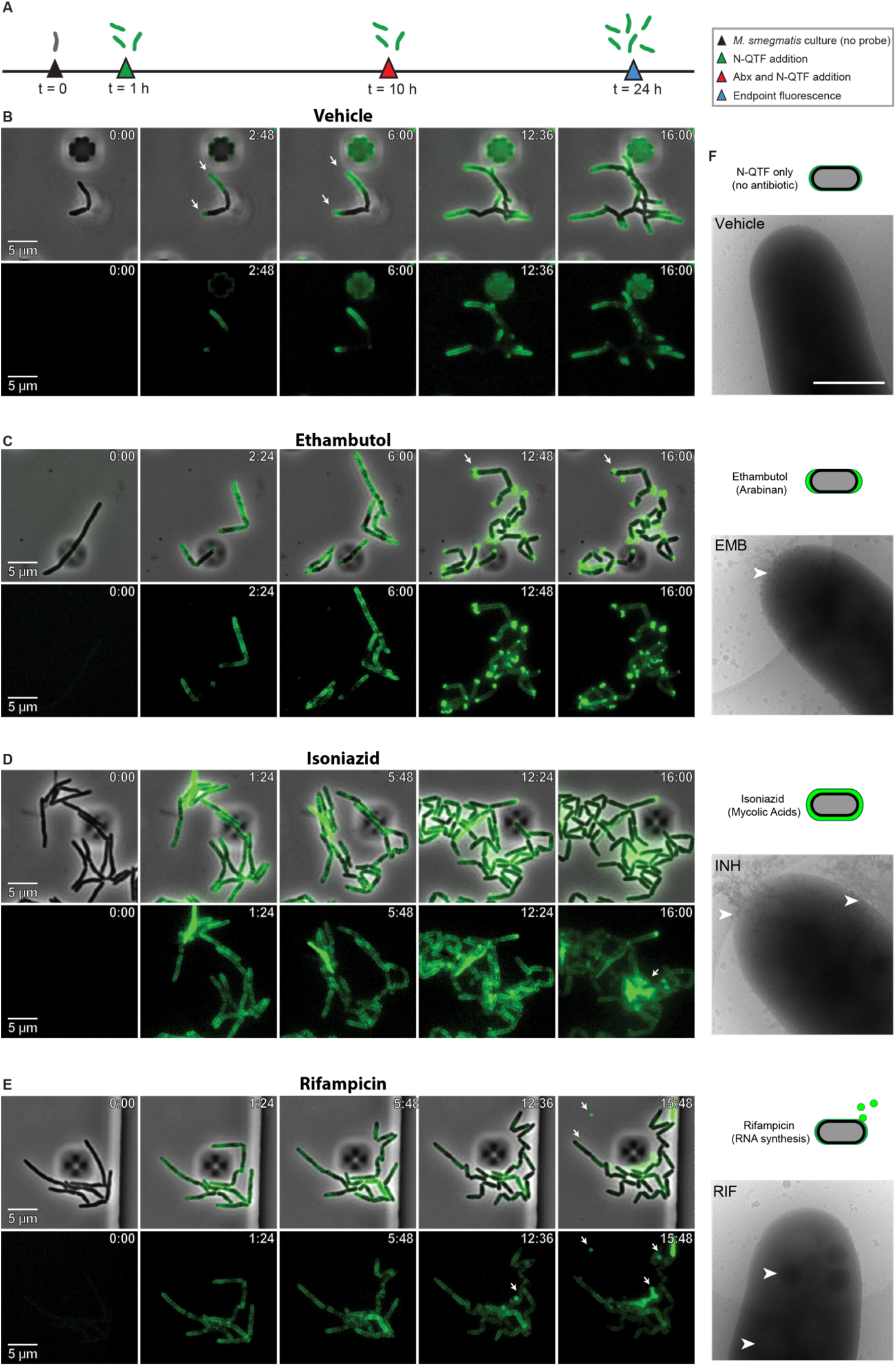
Antibiotic-specific phenotypes from fluorescence and superimposed phase imaging of *Msmeg*. (A) N-QTF and antibiotic treatment timeline used to monitor antibiotic phenotypes. (B) Combined microfluidics and time-lapse microscopy were used to assess antibiotic phenotypes relative to vehicle control following (C) EMB treatment, (D) INH treatment, (E) RIF treatment. (F) Cryo-transmission electron microscopy after four hours of exposure. Arrows indicate antibiotic phenotypes. Images are presented for vehicle, or treatment with EMB, INH, or RIF. Data are representative of two individual experiments.

We reasoned that other cell wall biosynthesis inhibitors might exhibit similar lipid localization phenotypes. The bactericidal frontline antibiotic INH targets the enoyl reductase InhA involved in cell envelope construction and perturbs the available mycolic acid lipid pool. Time-lapse imaging revealed that a delayed fluorogenic phenotype after one doubling time (Fig 2D). The fluorescence was dispersed throughout the cell membrane, supporting the known lytic mechanism of action of INH at a super-MIC concentration (25). Cell surface blebbing followed by membrane shredding has been reported to precede lytic activity in INH-treated *Msmeg* (26). Unexpectedly, we observed that EMB and INH, which both target cell envelope biosynthesis, did not produce similar phenotypes, suggesting differential drug mechanism of action results in altered lipid localization phenotypes in mycobacterial membranes.

We next tested whether antibiotics that block processes that do not directly impact cell envelope biogenesis would afford mycomembrane perturbations. To this end, we treated *Msmeg* with RIF, an antibiotic that engages bacterial RNA polymerase and blocks the initiation of RNA synthesis. Time-lapse imaging revealed the RIF treatment promoted the appearance of outer membrane vesicles (OMVs) (Fig. 2E). This phenotype was absent in either EMB- and INH-treated samples, indicating a RIF-specific stress response. Bacteria, including *Mtb*, actively release outer membrane vesicles in the absence of stress (16, 27, 28); however, enhanced mycobacterial OMV production following antibiotic treatment had not been previously observed. Imaging of N-QTF in RIF treated cells reveals previously uncharacterized cell wall activity upon inhibition of bacterial transcription.

### Cryo-TEM of antibiotic-treated mycobacteria

Guided by our dynamic phenotypic profiling, we used cryo-transmission electron microscopy (cryo-TEM) to gather additional insight into the varied antibiotic phenotypes. To this end, we treated *Msmeg* with EMB, INH, or RIF at sublethal doses. After four hours of antibiotic exposure, cryo-TEM images revealed dramatic differences analogous to those observed in time-lapse microscopy (Fig. 2F). EMB-treated bacteria showed lipid droplet accumulation at the cell poles. INH treatment caused lipid accumulation along the cell length and polar regions. RIF addition elicited the production of phosphate and lipid-rich subcellular bodies (29) throughout the bacillus, suggesting a priming step occurs before OMV release. These data indicate that N-QTF does not contribute to the observed phenotypes but strictly reports on cell remodeling.

### Rif-induced OMVs modulate host macrophage responses

The distinct OMV phenotype observed in RIF-treated mycobacteria was intriguing. Cryo-TEM could not capture this time-dependent phenotype, and previous studies indicate that secreted mycobacterial OMVs can modulate host immunity through TLR agonism (16). In this way, OMVs can help *Mtb* subvert immune surveillance by macrophages (16). We posited that the OMV phenotype observed in RIF-treated mycobacteria might differ from OMVs produced by *Mtb* in the absence of stress, thus providing insight into the recent rise in *Mtb* resistance to RIF (30). Indeed, RIF-treated infected macrophages can give rise to RIF-resistant *Mtb* with modified lipid envelopes (31). To query how mycobacterial OMVs under antibiotic treatment modulate host immune response, we isolated OMVs from the supernatant of DMSO vehicle-, EMB-treated, or RIF-treated *Mtb*. and used them to stimulate human blood monocyte-derived macrophages (Fig. 3A). We compared the responses to that of lipopolysaccharide from Gram-negative *Escherichia coli*, a potent stimulant of the human immune system. RT-qPCR analysis of primary human macrophages incubated with OMVs from DMSO-treated *Mtb* afforded an increase in the expression of primary proinflammatory cytokines. Notably, treatment of primary human macrophages with OMVs from RIF-treated *Mtb* decreased proinflammatory cytokine expression relative to OMVs from vehicle-treated *Mtb*. To further explore how a range of frontline antibiotics influence OMV-mediated immune modulation, we isolated OMVs from EMB-treated *Mtb*. The proinflammatory cytokine response from EMB OMVs was like that of the DMSO OMV control across the proinflammatory cytokines IL-1b, IL-6, and IL12b (Fig 3C). These data indicate that the distinct phenotypes observed in our imaging studies have different consequences on immune activation. The data also suggest that cell envelope remodeling by *Mtb* in response to RIF treatment can help evade host defense.

**Figure 3.**
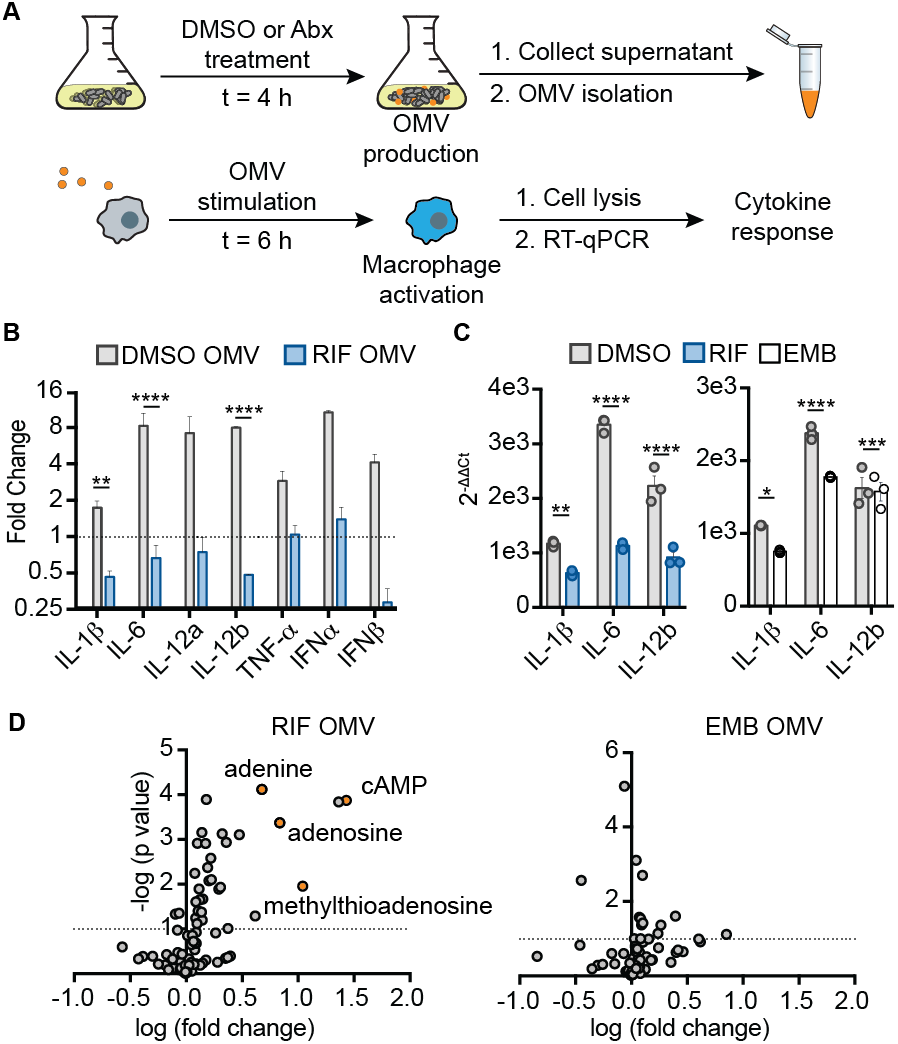
Assessment of human macrophage responses to OMVs from *Mtb*. (A) Schematic depiction of experimental workflow. OMVs were isolated from *Mtb* following treatment with DMSO, RIF, or EMB and exposed to human macrophages. Cytokine expression was measured following macrophage stimulation with purified OMVs. (B) RT-qPCR analysis of macrophage cytokine expression in response to RIF OMVs relative to LPS control. (C) A comparison of macrophage cytokine expression elicited by RIF OMVs compared to that from EMB OMVs. (Two-way ANOVA ****p ≤ 0.0001, ***p ≤ 0.001, **p ≤ 0.01, *p ≤0.05 and non-significant >0.05). (D) Targeted metabolomics of polar metabolites present in OMVs represented as log fold change of the average metabolite in RIF-or EMB-treated *Mtb* over DMSO-treated control. Data are representative of two independent experiments.

OMVs help *Mtb* and other bacteria deliver biomolecules at a high effective concentration. To explore the molecular basis of macrophage modulation by antibiotic-treated OMVs, we performed a targeted metabolomics analysis of 167 metabolites (Fig. 3D) (32). We detected that metabolites in the adenosine pathway were upregulated in RIF OMVs over DMSO OMVs. The largest fold-change was observed for cyclic AMP (cAMP), suggesting a link between intra-membrane vesicle cyclic AMP concentration and immune modulation. Since EMB OMVs did not significantly activate the proinflammatory macrophage response, we hypothesized that cAMP would not be present in EMB OMV metabolites. Consistent with this hypothesis, targeted metabolomics analysis of EMB OMVs showed no increase in cAMP or other adenosine metabolites (Fig. 3D). These data highlight that *Mtb*-derived OMVs can deliver nucleotide derivatives.

## Discussion

The glycolipid trehalose dimycolate is the major constituent of the mAG and confers protection against environmental stress. Unlike the arabinogalactan layer, the outermost mycolic acid layer is noncovalently assembled. Thus, we reasoned antibiotic-induced spatiotemporal rearrangement of the mycomembrane could be monitored with a glycolipid probe. To this end, we designed a stable, fluorogenic reporter of the mycobacterial membrane, N-QTF, to reveal dynamic cellular phenotypes critical to the bacterial life cycle. The simultaneous monitoring of membrane assembly and drug response was unprecedented and revealed insights into antibiotic-induced immunomodulatory mechanisms. Therefore, N-QTF affords new opportunities to identify concealed mechanisms of mycobacterial immunomodulation and discover new antibiotics to combat mycobacterial antibiotic resistance.

Genetic knockouts (33) used to identify essential gene targets are lethal by definition and fail to report on responsive mechanisms critical to mycobacterial survival. Emerging genetic (34–36) and morphological (12) strategies have revealed mycobacterial vulnerabilities and unique mechanisms of antibiotic action, but lack information of dynamic remodeling phenotypes in active drug response mechanisms. N-QTF’s unique ability to report on active remodeling and secretion mechanisms was critical to decoding the immunomodulatory role of antibiotic-induced secretion of OMVs from *Mtb*.

Our finding of cAMP’s presence in antibiotic-induced OMVs was unanticipated. Changes in intramacrophage cAMP levels have been observed in *Mtb* infection; however, a role for antibiotics in altering cAMP levels was unknown. Increases in cAMP can dampen macrophage signaling through protein kinase A and the cAMP response element binding protein pathways (37–39). Moreover, adenosine and methylthioadenosine have been shown to reduce the secretion of the proinflammatory cytokine TNF-α in macrophages by attenuating Toll-like receptor activation pathways (40). The delivery of high local concentrations of encapsulated adenosine metabolites in response to RIF may allow *Mtb* to control host macrophage immune responses, suggesting a mechanism for bacterial survival and subsequent host tolerance.

The mycobacterial membrane grants *Mtb* recalcitrance to antimicrobials. How the mycomembrane dynamically remodels in response to antibiotic stress was poorly understood. By leveraging a substrate mimic of the enzymes that control mycomembrane remodeling, we investigated changes in membrane phenotypes from frontline antibiotic treatment without genetic manipulation. The critical observation that the spatial and temporal localization of the lipids is dependent on an antibiotic’s mechanism of action highlights the utility of our approach. The secretion of OMVs from RIF-treated mycobacteria was unprecedented and suggests these bacteria modulate the host immune environment as collateral damage to their stress response (Supp. Fig. S5). Our findings indicate that consideration of the consequences of bacterial treatment on host immune responses could yield more effective strategies for antibiotic targeting of *Mtb*. Last, we demonstrate how agents that report on the mycomembrane can be leveraged to report on the biology of existing antibiotics and discover new mechanisms of action.

## Materials and Methods

### General Information for Synthetic Methods and Materials

All reagents for synthetic starting materials, buffers, and antibiotics were used as received from commercial manufacturers unless otherwise noted. Organic solvents were purified. Water was purified using a Millipore Milli-Q Integral Water Purification system. All reactions were conducted under an inert nitrogen atmosphere in oven-dried glassware unless otherwise stated. Analytical thin-layer chromatography (TLC) was carried out on E. Merck (Darmstadt) TLC plates pre-coated with silica gel 60 F254 (250 μm layer thickness). Analyte visualization was accomplished using a UV lamp or by charring with a phosphomolybdic acid solution. Flash column chromatography was performed with Silicycle flash silica gel (40−63 μm, 60 Å pore size) using the quoted eluent. 1H and 13C nuclear magnetic resonance (NMR) spectra were recorded on a 400 MHz spectrometer (acquired at 400 MHz for 1H NMR and 101 MHz for 13C NMR), or a 500 MHz spectrometer (acquired at 500 MHz for 1H NMR and 126 MHz for 13C NMR). NMR chemical shifts are reported as follows: chemical shift (δ ppm), multiplicity (s = singlet, d = doublet, t = triplet, q = quartet, p = quintet, or some combination thereof. Chemical shifts are reported relative to residual solvent peaks in parts per million (CDCl3: 1H, 7.27, 13C, 77.23; CD3OD: 1H, 3.31, 13C, 49.15). Coupling constants (J) are reported in Hertz (Hz) and rounded to the nearest 0.1 Hz. High-resolution mass spectra (HRMS) were obtained on an electrospray ionization-time of flight (ESI-TOF) Micromass LCT mass spectrometer.

### Bacterial strain selection and culture conditions

Mycobacterial species *M. smegmatis* mc^2^155, *M. marinum* BAA-535, and *M. tuberculosis* H37Rv were grown with shaking at 37 °C in Middlebrook 7H9 broth (BD, Franklin Lake, NJ) with 0.2% (v/v) glycerol, 0.05% (v/v) Tween-80 with or without OADC supplement (Sigma-Aldrich).

### Time-dependent fluorescence assays

Mycobacterial species *M. smegmatis* mc^2^155, *M. marinum* BAA-535, and *M. tuberculosis* H37Rv were grown to OD600 = 1. Strains were diluted to OD600=0.2. Reactions were initiated by the addition of N-QTF to a final volume of 50 μL per well (n=3). Relative fluorescence units were measured at ex/em 485/525 and read every 15 minutes for two doubling times (t=6 hours) at 37 °C.

### Purification of *Mtb*Ag85 mycolyltransferases

*Mtb* Ag85A, Ag85B, and Ag85C were obtained through BEI Resources, NIAID, NIH: Purified native protein from *M. tb*. H37Rv antigen 85A (fbpA, Rv3804c, cat. No: NR-14856), antigen 85B (fbpB, Rv1886c, cat. No: NR-14857), and antigen 85C (fbpD, Rv3803, cat no: NR-14858). Upon receipt, lyophilized native Ag85 proteins were thawed and suspended in aqueous 1xPBS buffer (Gibco cat no:10010023) to a concentration of 10 μM for storage at –20 °C. *Mtb* Ag85A–C proteins were thawed and used as needed.

### Kinetic assays with *Mtb* Ag85A-C mycolyltransferases

A flat, clear-bottom 96-well black polystyrene microplate (Greiner, cat no: 655097) was used to measure time-dependent N-QTF activation with 5 μM N-QTF with a fixed concentration (1 μM) of each Ag85 isoform in 1x PBS pH 7.4. Reactions were initiated by the addition of N-QTF to a final volume of 50 μL per well (n=3). Relative fluorescence units were measured at ex/em 485/525 and read every 30 seconds for one hour at 37 °C.

### Fluorescence microscopy

Mycobacterial species *M. smegmatis* mc^2^155, *M. marinum* BAA-535, and *M. tuberculosis* H37Rv were grown to OD600 = 1 in either Middlebrook 7H9 or 7H9 +OADC supplement 0.2% glycerol, 0.05% Tween-80. Strains were diluted to OD600=0.2 and N-QTF was added at a final concentration of 2.5 uM in 1 mL of liquid culture. 10 uL of culture was removed after 24 hours and placed on a glass microscope slide with polylysine-coated coverslips. Slips were sealed and imaged on a Cell Discoverer7 at 100x magnification. Visualization of N-QTF was performed and monitored at ex/em 488/525 nm. Imaging processing was performed using ImageJ FIJI software.

### Antibiotic-dependent N-QTF incorporation

A flat, clear-bottom 96-well black polystyrene microplate (Greiner, cat no: 655097) was used to measure time-dependent N-QTF (1 μM) activation with frontline antibiotic treatment to *M. smegmatis* mc2155. EMB, INH, or RIF was used to treat *M. smegmatis* at OD600 = 0.2 at either 50 μg/mL or 12.5 μg/mL (n=3). Relative fluorescence units were measured at ex/em 485/525 and read every 15 minutes for two doubling times (t=6 hours) at 37 °C.

### Time-lapse microscopy

*M. smegmatis* were grown to mid-log and diluted to OD600 of 0.3. Cells were loaded into a CellASIC (cat no: B04A) plate under constant microfluidic flow with 7H9 medium in a 37 °C chamber and allowed to grow in the chamber for one hour. 7H9 medium supplemented with N-QTF (1 μM) was flowed into the chamber for six hours before switching media to N-QTF with and without antibiotic (10 μg/mL). Images were acquired every 15 min on an inverted Nikon TI-E microscope using a 60× objective. Cells were imaged using phase contrast and ex/em at 488/525 nm.

### Cryo-transmission electron microscopy

Cells were cultured to mid-log in complete 7H9 with and without frontline antibiotic treatment at 10 μg/mL for four hours at 37 °C. After four hours, cells were concentrated 50-fold by pelleting by centrifugation and resuspending in a 50-fold smaller volume. Concentrated cells were plunge-frozen in liquid ethane using an automated Leica EM GP system (Leica Microsystems) using R2/2 200 mesh grids (Quantifoil) and imaged using a Talos 120 kV cryo-transmission electron microscope.

### Monocyte-derived macrophage differentiation and purification

Adult human peripheral blood was acquired from Research Blood Components, LLC (Boston, MA, USA) under IRB-approved protocol. Monocytes were isolated from whole blood by negative selection using RosetteSep human monocyte enrichment cocktail following the manufacturer’s protocol (StemCell Technologies). Using density gradient medium (Lymphoprep) monocytes were isolated and differentiated into macrophages in the presence of GM-CSF (100 ng/mL) (R&D Systems) in 1640 RPMI + 5% FBS +P/S media for 6 to 7 days

### Outermembrane vescicle isolation

*Mtb* H37Rv (500 mL) cultures were treated with DMSO, EMB, or RIF at 15 ug/mL final concentration for six hours at 37 °C. The supernatant was centrifuged at 38,000xg in a JA16.25 (Beckman Coulter) for 3 hours at 4 °C. After centrifugation, the supernatant was removed and the pellet was resuspended in 10 mM HEPES 150 mM NaCl pH 7.2. OMV concentration was normalized using absorbance at 280 nm and aliquoted into 100 μL samples and stored at -20 °C.

### OMV stimulation assay and Quantitative Real-Time PCR

Vehicle, EMB, or RIF OMVs were added to wells containing 500 μL of 750,000 cells/mL and incubated at 37 °C for 6 hours. After 6 hours, the supernatant was removed and washed with 2x Cold 1xPBS (1 mL). Trizol reagent (400 μL) was added to the stimulation well. The supernatant was removed at 300 μL of trizol was added. Total RNA was extracted with Trizol using Direct-zol RNA kits (Zymo Research). cDNA was generated by using an iScript cDNA synthesis kit (BioRad), and PCR amplification was performed in the presence of SYBR Green (BioRad) in a CFX96 real-time PCR detection system (BioRad). Specific primers were designed using the PrimerQuest tool (Integrated DNA Technologies, Inc., Supplementary Table 1). Expression of specific genes was normalized to GAPDH expression (ΔCt), and expression fold change was calculated using the delta Ct method (2−(ΔCt Stim-ΔCt Unstim)) and normalized to LPS (50 μg).

### Polar metabolite isolation

OMV samples from DMSO, EMB, and RIF-treated conditions were extracted to analyze the metabolite composition within them. In a microcentrifuge tube, OMVs (100 μL, n=3) were resuspended in 300 μL of LC/MS grade methanol containing a mixture of 17 isotope-labeled amino acids (200 nM, Cambridge Isotope Laboratories, MSK-A2-1.2). To this solution, LC/MS grade water (150 μL) was added to each tube. Ice cold amylenes-free chloroform (200 μL) was added to each sample. The sample was vortexed at 4 °C for 1 minute and centrifuged for 10 minutes at 13,000xg at 4 °C. The top aqueous layer was removed and allowed to dry under N_2_ stream overnight. Dried extracts were stored at -80 ºC until run.

### Targeted metabolomics

LC/MS was used to profile and quantify the polar metabolite contents of OMVs from DMSO, EMB, or RIF-treated *M*.*tb*.LC/MS-based analyses were performed as described previously(41) on a QExactive benchtop orbitrap mass spectrometer equipped with an Ion Max source and a HESI II probe coupled to a Dionex UltiMate 3000 UPLC system (Thermo Fisher Scientific). Metabolite sample (5 μL) was injected onto a ZIC-pHILIC 2.1 × 150 mm (5 μm particle size) column (EMD Millipore). Buffer A was 20 mM ammonium carbonate, 0.1% ammonium hydroxide; buffer B was acetonitrile. A flow rate of 0.150 ml/min was used for the chromatographic gradient. 0–20 min: linear gradient from 80% to 20% B; 20–20.5 min: linear gradient from 20% to 80% B; 20.5–28 min: hold at 80% B. The mass spectrometer was operated in full-scan, polarity switching mode with the spray voltage set to 3.0 kV, the heated capillary held at 275 °C, and the HESI probe held at 350 °C. The sheath gas flow was set to 40 units, the auxiliary gas flow was set to 15 units, and the sweep gas flow was set to 1 unit. The MS data acquisition was performed in a range of 70–1000 m/z, with the resolution set at 70,000, the AGC target at 10^6^, and the maximum injection time at 80 ms. XCalibur QuanBrowser 2.2 (Thermo Fisher Scientific) was used for metabolite identification and quantification using a 10 ppm mass accuracy window and 0.5 min retention time window compared to authentic metabolite standards. Within-batch mass deviation was typically < 0.0005 Da, and retention time deviation was < 0.25 min. In each sample, the raw peak area for each metabolite was divided by the raw peak area of the relevant isotope-labeled internal standard to calculate the relative abundance.

### Detailed synthetic procedures

#### Synthesis of 6-O-(15-[BODIPY-FL]-pentadeconyl)-6’-NH-DABCYL- α,α-trehalose/N-QTF 1 (See Figure S1 for overall route)

Dowex-50WX8-200 ion exchange resin (32 mg) was added to a solution of compound 5 (7.1 mg, 0.0044 mmol) in methanol (1.25 mL) and stirred at rt for 1 hr. The resin was removed by filtration and washed with methanol. The filtrate was concentrated under reduced pressure. Purification of the resulting residue by column chromatography (SiO_2_, 10→15% MeOH/CH_2_Cl_2_) afforded the title compound 1 (3.24 mg, 62%) as an orange solid: R_*f*_ = 0.20 (10% MeOH/CH_2_Cl_2_); ^1^H NMR (500 MHz, CD_3_OD) δ 7.97 – 7.95 (m, 2H), 7.88 – 7.85 (m, 4H), 7.74 (s, 1H), 7.42 (s, 1H), 6.98 (d, J = 4.0 Hz, 1H), 6.86 – 6.84 (m, 2H), 6.30 (d, J = 4.0 Hz, 1H), 6.22 (s, 1H), 5.10 (t, J = 4.1 Hz, 2H), 4.61 (dd, J = 11.9, 2.1 Hz, 1H), 4.44 (dd, J = 11.8, 5.4 Hz, 1H), 4.41 (s, 2H), 4.39 (dd, J = 11.9, 2.2 Hz, 1H), 4.32 (t, J = 7.1 Hz, 2H), 4.22 – 4.19 (m, 2H), 4.03 (ddd, J = 10.2, 5.5, 2.2 Hz, 1H), 3.82 (ddd, J = 15.1, 9.7, 8.9 Hz, 2H), 3.54 – 3.47 (m, 3H), 3.33 (dd, J = 10.1, 8.9 Hz, 2H), 3.25 (t, J = 7.6 Hz, 2H), 3.12 (s, 6H), 2.66 (t, J = 7.6 Hz, 2H), 2.52 (s, 3H), 2.34 (t, J = 7.4 Hz, 2H), 2.28 (s, 3H), 1.84 (p, J = 7.1 Hz, 2H), 1.61 (p, J = 7.3 Hz, 2H), 1.31 – 1.23 (m, 20H) ppm; ^13^C NMR (126 MHz, CD3OD) δ 174.1, 173.2, 160.0, 155.1, 153.3, 144.9, 144.4, 143.4, 135.2, 134.3, 133.5, 128.1, 128.0, 125.0, 124.4, 122.7, 121.6, 120.0, 116.3, 111.2, 94.0, 93.9, 73.2, 72.9, 72.1, 71.9, 71.8, 70.8, 70.4, 70.2, 62.9, 49.9, 48.2, 48.0, 47.9, 47.2, 47.1, 40.8, 39.0, 34.3, 33.7, 29.9, 29.3, 29.2, 29.1, 28.9, 28.7, 28.7, 26.0, 24.7, 13.5, 9.8 ppm; HRMS (ESI-TOF+) calcd for C_59_H_81_BF_2_N_10_O_13_ (M+H+) 1186.6155, found 1187.6128.

#### Synthesis of 6’-NH-DABCYL-2,3,4,2’,3’,4’-hexakis-O-(trimethylsilyl)-α,α-trehalose 3

DABCYL-OH (62 mg, 0.23 mmol), EDC•HCl (51mg, 0.26 mmol), DMAP (2.7 mg, 0.02 mmol), and HOBt (41 mg, 0.26) were dissolved in dry CH_2_Cl_2_ (1 mL) and stirred at rt for 15 min, then cooled to 0 °C. A solution of compound **2** (187 mg, 0.22 mmol) in dry CH_2_Cl_2_ (6 mL) was added dropwise via cannula, and the reaction mixture kept at 0 °C with slow addition of DIPEA (192 μL). Reaction was allowed to warm to rt for 20 h and then concentrated under reduced pressure and resuspended in CH_2_Cl_2_ (20 mL) and a cooled solution of 0.2 M Na_2_CO_3_/NaHCO_3_ buffer pH=9 (20 mL). The crude reaction mixture was immediately transferred to a separatory funnel extracted with CH_2_Cl_2_, washed with brine, and dried over Na_2_SO_4_ to afford crude material. Purification of the resulting residue by column chromatography (SiO_2_, 0→8% EtOAc/ CH_2_Cl_2_) afforded the title compound **3** (150 mg, 66%) as an orange solid: R_*f*_ = 0.35 (30% EtOAc/hexanes); ^1^H NMR (400 MHz, CDCl_3_) δ 7.75 – 7.73 (m, 2H), 7.71 – 7.73 (m, 4H), 6.61 – 6.60 (m, 2H), 6.30 (d, J = 3.1 Hz, 2H), 4.81 (d, J = 2.4 Hz, 1H), 4.75 (d, J = 3.7 Hz, 1H), 3.82 (dq, J = 8.4, 2.7 Hz, 1H), 3.80 (t, J = 8.9 Hz, 1H), 3.78 (t, J = 8.9 Hz, 1H), 3.75 (dt, J = 9.5, 3.4 Hz, 1H), 3.33 – 3.29 (m, 3H), 3.26 – 3.20 (m, 3H), 2.95 (s, 6H), 1.42 (dd, J = 7.4, 5.3 Hz, 1H), 0.04 (s, 9H), 0.00 (s, 9H), 0.00 (s, 9H), - 0.01 (s, 9H), -0.01 (s, 9H), -0.11 (s, 9H) ppm; ^13^C NMR (101 MHz, CDCl3) d 166.7, 154.9, 152.7, 143.5, 134.4, 127.7, 125.3, 122.1, 111.4, 94.2, 93.8, 73.9, 73.2, 73.1, 73.0, 72.9, 72.7, 72.6, 71.3, 61.6, 41.4, 40.2, 0.94, 0.91, 0.75, 0.23, 0.00, 0.00 ppm; HRMS (ESI-TOF+) calcd for C_45_H_84_N_4_O_11_Si_6_ (M+H+) 1025.4825, found 1025.44830.

#### Synthesis of 6-O-(15-azidopentadeconyl)-6’-NH-DABCYL-2,3,4,2’,3’,4’-hexakis-O-(trimethylsilyl)-α,α-trehalose 4

15-Azidopentadecanoic acid (23 mg, 0.082 mmol), EDC•HCl (31 mg, 0.164 mmol) and DMAP (6 mg, 0.049 mmol) were dissolved in dry CH_2_Cl_2_ (0.5 mL) and stirred at rt for 15 min. A solution of compound **3** (42 mg, 0.041 mmol) in dry CH_2_Cl_2_ (1 mL) was added dropwise via cannula, and the reaction mixture stirred at rt for 16 h and then concentrated under reduced pressure. Purification of the resulting residue by column chromatography (SiO_2_, 0→4% EtOAc/hexanes) afforded the title compound **4** (36 mg, 69%) as an orange solid: R_*f*_ = 0.33 (10% EtOAc/hexanes); ^1^H NMR (400 MHz, CDCl_3_) δ 7.74 (m, 2H), 7.72 – 7.67 (m, 4H), 6.61 – 6.58 (m, 2H), 5.13 (t, J = 3.6 Hz, 2H), 4.82 (dd, J = 12.0, 2.4 Hz, 1H), 4.17 (ddd, J = 11.2, 7.9, 2.8 Hz, 2H), 3.91 – 3.72 (m, 5H), 3.31 (t, J = 9.0 Hz, 1H), 3.27 – 3.21 (m, 3H), 3.08 (t, J = 7.0 Hz, 2H), 2.94 (s, 6H), 2.19 (td, J = 7.5, 3.5 Hz, 2H), 1.48 – 1.39(m, 4H), 1.20 – 1.08 (m, 20H), 0.04 (s, 9H), 0.00 (s, 9H), -0.01 (s, 9H), -0.01 (s, 9H), -0.03 (s, 9H), -0.12 (s, 9H) ppm; ^13^C NMR (101 MHz, CDCl3) δ 172.7, 165.7, 154.0, 151.7, 142.6, 133.5, 133.4, 126.8, 124.3, 121.2, 110.4, 72.9, 72.5, 72.3, 72.2, 72.0, 71.8, 71.7, 71.5, 70.8, 70.4, 70.1, 69.8, 62.2, 61.3, 52.4, 50.4, 40.5, 39.2, 33.1, 28.6, 28.5, 28.4, 28.4, 28.3, 28.1, 28.1, 27.8, 25.6, 23.7 0.00, 0.00, 0.00, -0.18, -0.88, -0.94 ppm; HRMS (ESI-TOF+) calcd for C_60_H_111_N_7_O_12_Si_6_ (M+H+) 1290.6979, found 1290.6980

#### Synthesis of 6-O-(15-[BODIPY-FL]-pentadeconyl)-6’-NH-DABCYL-2,3,4,2’,3’,4’-hexakis-O-(trimethylsilyl)-α,α-trehalose 5

Compound **4** (12 mg, 0.009 mmol) and BODIPY-FL alkyne (3 mg, 0.009 mmol) were dissolved in 5:5:1 t-BuOH:H_2_O:CH_2_Cl_2_ (1.5 mL total). To this solution were added TBTA (97 μg in 96.7 μL CH_2_Cl_2_, 0.0018 mmol), sodium ascorbate (18 μL of a 0.1M aqueous solution, 0.0018 mmol), and CuSO_4_•5H_2_O (9 μL of a 0.1M aqueous solution, 0.0009 mmol). The resulting mixture was stirred vigorously in the dark for 48 h, and then concentrated under reduced pressure. Purification of the resulting residue by column chromatography (SiO2, 40→90% EtOAc/hexanes) afforded the title compound **5** (11.2 mg, 76%) as an orange solid: R_*f*_ = 0.15 (50% EtOAc/hexanes); ^1^H NMR (500 MHz, CDCl_3_) δ 7.74 – 7.67 (m, 2H), 7.70 – 7.67 (m, 4H), 7.10 (s, 1H), 6.90 (s, 1H), 6.68 (d, J = 4.0 Hz, 1H), 6.60 (d, J= 4 Hz, 1 H), 6.31 – 6.29 (m, 2H), 6.07 (m, 1H), 5.95 (s, 1H), 4.82 (dd, J = 5.2, 3.1 Hz, 2H), 4.75 (dd, J = 11.9, 2.4 Hz, 1H), 4.32 (d, J = 5.8 Hz, 2H), 4.17 – 4.09 (m, 4H), 3.96 (dt, J = 9.5, 3.0 Hz, 1H), 3.90 (dd, J = 11.8, 4.5 Hz, 1H), 3.88 (ddd, J = 9.6, 4.5, 2.3 Hz, 1H), 3.81 (t, J = 8.9 Hz, 1H), 3.80 (t, J = 8.9 Hz, 1H), 3.75 (t, J = 9.1 S4 Hz, 1H), 3.33– 3.23 (m, 3H), 3.22 (t, J = 7.5 Hz, 2H), 2.94 (s, 6H), 2.48 (t, J = 7.6 Hz, 2H), 2.39 (s, 3H), 2.23 – 2.15 (m, 2H), 2.08 (s, 3H), 1.69 (q, J = 7.3 Hz, 2H), 1.46 – 1.43 (m, 4H), 1.13 – 1.07 (m, 18H), 0.04 (s, 9H), 0.00 (s, 9H), -0.01 (s, 9H), -0.01 (s, 9H), 0.03 (s, 9H), -0.12 (s, 9H) ppm; ^13^C NMR (126 MHz, CDCl_3_) δ 172.7, 170.2, 165.7, 159.4, 156.0, 154.0, 151.8, 142.9.9, 142.6, 134.2, 133.4, 132.3, 127.0, 126.8, 124.4, 122.7, 121.2, 121.0, 119.5, 116.2, 110.4, 93.1, 92.7, 72.9, 72.0, 71.8, 71.5, 70.8, 70.3, 69.8, 62.2, 49.3, 40.5, 39.3, 34.8, 34.0, 33.1, 29.2, 28.7, 28.6, 28.5, 28.4, 28.4, 28.3, 28.1, 28.0, 25.5, 23.8, 13.9, 13.2, 13.1, 10.3, 0.2, 0.0, -0.2, -0.2, -0.9, -0.9; HRMS (ESI-TOF+) calcd for C_77_H_129_BF_2_N_10_O_13_Si_6_ [(M+2H+)/2] 810.4281, found 810.428.

## Acknowledgments

The authors thank Dr. Eachan Johnson, Dr. Amanda Dugan, and Dr. Sharon Wong for helpful discussions. Additional thanks to Dr. Caroline Lewis, Tenzin Kunchok, and Dr. Kayla Crowder of the Metabolite Profiling Core Facility at the Whitehead Institute for metabolomics analysis and data. This researcher was supported by the National Institute of Allergy and Infectious Disease (R01-AI126592 to L.L.K.; R21-AI156772, to E.J.R.). MGW was funded by the Broad Institute TB Gift Innovation Award and the NIH Ruth L. Kirschstein National Research Service Award (F32-GM133116).

